# *In silico* Identification of Novel Common Drug Targets Against Four Infectious *Acinetobacter* Species

**DOI:** 10.1101/2024.11.07.622554

**Authors:** Soaibur Rahman, Soharth Hasnat, M Murshida Mahbub, Zinat Farzana, Sal Sabila

**Author notes:** **Correspondence to:** M Murshida Mahbub.

## Abstract

The global emergence of multidrug-resistant *Acinetobacter* species has become a major concern in the management of hospital-acquired infections. Moreover, misdiagnosis of one species of *Acinetobacter* with another has been reported, making it difficult to choose appropriate treatment. World Health Organization emphasizes the urgent need to develop new antibiotics to combat *Acinetobacter* infections. This study aimed to discover novel common drug targets that will be effective against *Acinetobacter nosocomialis, Acinetobacter baumannii, Acinetobacter pittii,* and *Acinetobacter haemolyticus*. We utilized a cluster-based subtractive genomics approach to identify potential drug targets. The main focus was to find drug targets that are absent in humans and essential for pathogens. We also performed metabolic pathway and subcellular localization analyses. Furthermore, protein structure-based studies and druggability analyses were conducted to identify viable therapeutic options. Out of 1245 protein clusters (minimum 4 proteins/cluster), 204 clusters were human non-homologous and essential for bacteria. Among them, 39 clusters were cytoplasmic and involved in unique metabolic pathways which are specific to the pathogens. After analyzing the drug target sequences of DrugBank database, 12 clusters were found to be novel drug targets. Eventually, proteins of one cluster were identified as advantageous drug targets having drug-binding pockets at very similar regions with high druggability scores. These proteins can be inhibited to disrupt the Lysine/DAP biosynthetic pathway of *Acinetobacter.* Our research might open up a new possibility for drug discovery against these pathogens.

## 1. Introduction

*Acinetobacter* species are strictly aerobic, gram-negative saprophytes that are found almost anywhere (Almasaudi, 2018). These species have become notorious nosocomial pathogens that affect immunocompromised patients in intensive care units (ICU) (Wisplinghoff et al., 2000). Hospital-acquired pneumonia, urinary tract infection, and bacteremia are some common infections caused by *Acinetobacter* spp. (Bergogne-Bérézin, 2008). Despite their low virulence, it is concerning that they have developed resistance against many common antibiotics such as ampicillin, tetracycline, gentamicin, ciprofloxacin, chloramphenicol, etc. (Almasaudi, 2018; Kiskova et al., 2023). CDC reported that approximately 8,500 infections occurred in hospitalized patients and around 700 died in the United States due to antibiotic-resistant *Acinetobacter* in 2017 (CDC, 2024).

The majority of infections and outbreaks in hospitals are caused by *Acinetobacter baumannii*, which accounts for 95% of all cases (Lupo et al., 2018). This opportunistic pathogen is responsible for around a million cases worldwide each year, with a high fatality rate (Cavallo et al., 2023). The capacity to live under harsh conditions, for example, nutrient-limited dry surfaces allows them to persist and spread in healthcare settings (Almasaudi, 2018). Multidrug-resistant *A*. *baumannii* has been identified in hospitals across continents (Perez et al., 2007). Although *Acinetobacter* species other than *A. baumannii* are often considered less dangerous, *A. nosocomialis* is equally infectious. *A. nosocomialis* infection is frequently confused with *A. baumannii* infection. It was responsible for 14 of the 48 cases of infection in Thailand that were initially thought to be caused by *A. baumannii* (Nithichanon et al., 2022). Another *Acinetobacter* species, *Acinetobacter pittii* is one of the members of *Acinetobacter calcoaceticus*-*Acinetobacter baumannii* (ACB) complex. A study investigated ACB complex bloodstream infections (BSI) in a French hospital and revealed that *A. pittii* was more frequently associated with BSI than *A. baumannii* (Pailhories et al., 2018). *A. Pittii* isolated from the harsh environment of the International Space Station (ISS) has shown more resistance against specific antibiotics than related clinical isolates (Tierney et al., 2022). It is currently evident that *A. Pittii* has a probability of accumulating more resistance genes, which could make it difficult to control non-*baumannii* ACB species in a medical setting (Pailhories et al., 2018). Likewise, *Acinetobacter haemolyticus* is another disease-causing species of *Acinetobacter*. Except for cefepime, aztreonam, and ceftazidime, *A. haemolyticus* demonstrated resistance to all β-lactams. The possibility of *A. haemolyticus* causing a nosocomial BSI outbreak in the future should be considered (Bai et al., 2020).

The emergence of pan-drug resistant *Acinetobacter* species necessitates the identification of novel therapeutic targets and the development of potential drugs. Currently, computational biology serves as a routine tool for identifying new compounds against specific drug targets (Hossain et al., 2024; Islam et al., 2024). In this study, however, we focus on target identification, applying a cluster-based subtractive genomics approach to uncover novel, conserved drug targets (Hasnat et al., 2024). Initially, core genomes of the four *Acinetobacter* species (namely, *Acinetobacter nosocomialis, Acinetobacter baumannii, Acinetobacter pittii, and Acinetobacter haemolyticus*) were identified by pan-genome analysis. Afterward, we ran several computational investigations and used multiple web servers to detect proteins that are non-human homologs, essential for pathogens, novel, and suitable as drug targets. We included structural and druggability analyses to further validate our findings.

## 2. Material and methods

In order to identify novel common drug targets against *Acinetobacter* nosocomialis, *Acinetobacter baumannii*, *Acinetobacter* pittii, and *Acinetobacter* haemolyticus, a cluster-based subtractive genomics approach was employed. Fig. 1 demonstrates the overall workflow in brief.

**Fig. 1.**
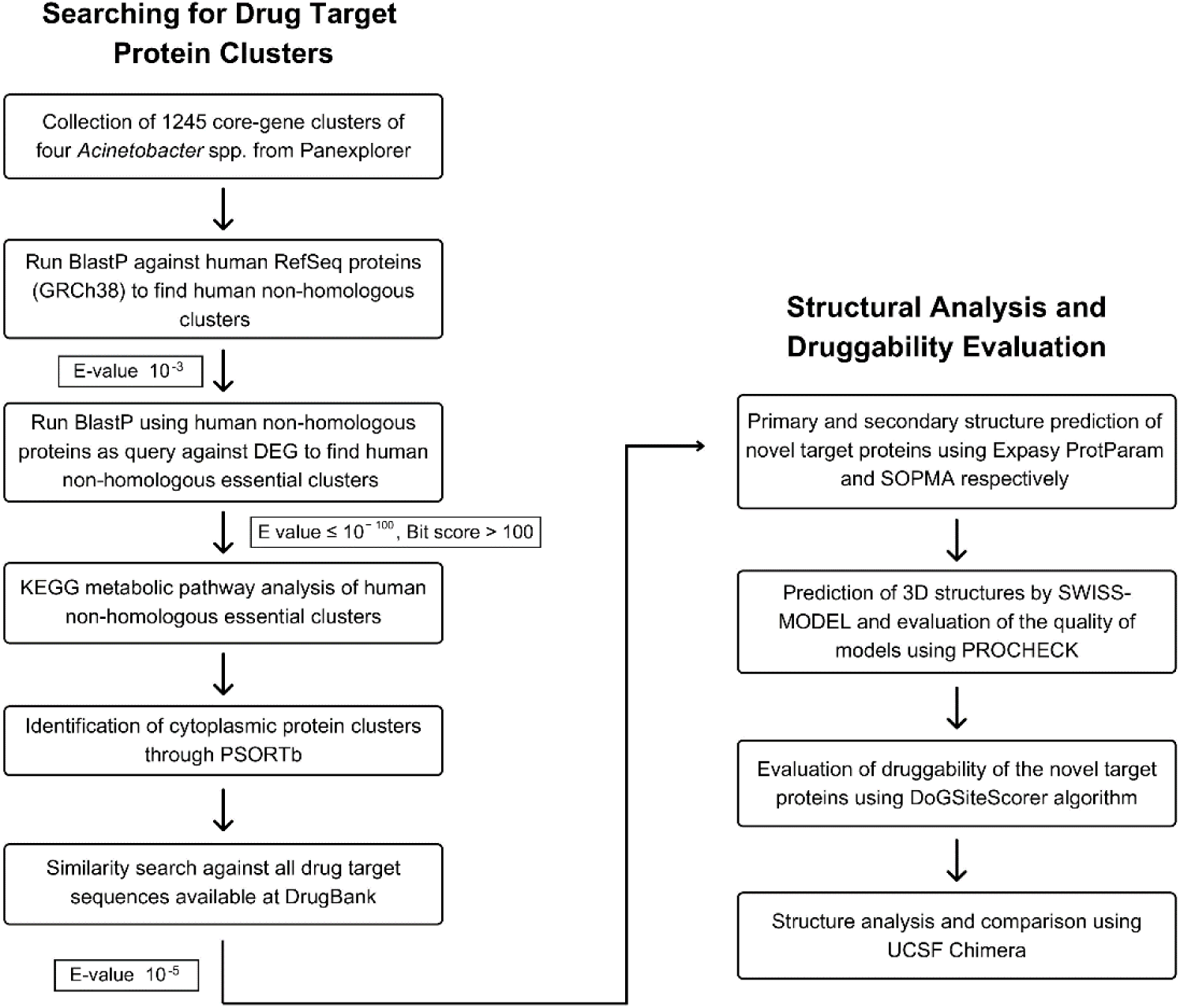
A step-by-step schematic representation of the methodology.

### 2.1. Collection of core-gene clusters

Genome accessions of *Acinetobacter baumannii* (AYE) (Genome accession: CU459141), *Acinetobacter haemolyticus* (TJS01) (Genome accession: CP018871), *Acinetobacter nosocomialis* (M2) (Genome accession: CP040105), and *Acinetobacter pittii* (PHEA-2) (Genome accession: CP002177) were collected from UniProt (UniProt, 2021) and submitted to Pan-genome Explorer (Hasnat et al., 2024). The minimum percent identity for BLAST was set at 80. Pan-genome Explorer is an online tool that utilizes PGAP pipeline to conduct pan-genome analysis and provides lists of gene clusters along with various data interpretations (Dereeper et al., 2022). A FASTA file was prepared that contained all the protein sequences of each core-gene cluster of the four *Acinetobacter* species.

### 2.2. Identification of human non-homologous clusters

BLASTp search for bacterial cluster sequences was performed against human RefSeq proteins (GRCh38) with a threshold of expect value (E value) 10^−3^ to identify bacterial clusters that are non-homologous to human proteome (Wadood et al., 2018). E value is commonly used for assessing the quality of sequence alignment. Lower E values denote more significant matches since they are less likely to be found just by chance. E value in BLASTp is calculated by the following formula:

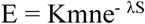

where the expect value is denoted by E. The lengths of the query sequence and the database are represented by m and n, respectively. The Karlin-Altschul parameters are K and λ. These statistical parameters depend on the scoring system and database composition. S refers to the alignment score (Altschul et al., 1990). This step is important to reduce the cross-reactivity of drugs with similar protein sequences in humans (Sarkar et al., 2012).

### 2.3. Identification of essential clusters

Human non-homologous protein cluster sequences were subjected to BLASTp against the Database of Essential Genes (DEG) (http://origin.tubic.org/deg/public/index.php/download) with E value ≤ 100^−100^ and Bit score > 100, where the bit score provides insight about the statistical significance of an alignment. Higher bit scores suggest a greater similarity between the sequences (Rahman et al., 2014). DEG is a collection of genes that are essential for cell survival. This database has emerged as a pivotal tool for essential gene focused studies. It compiles essential gene data acquired through various experimental approaches at a whole-genome scale (Liang et al., 2024; Zhang et al., 2004).

### 2.4. Analysis of metabolic pathways

Human non-homologous essential protein clusters were subjected to KAAS (KEGG Automatic Annotation Server). KAAS performs BLAST comparisons against the manually curated KEGG GENES database to give functional annotation of genes (Kanehisa and Goto, 2000). The result provided KO (KEGG Orthology) assignments and a list of metabolic pathways in which query sequences were involved. Human non-homologous essential clusters involved in unique metabolic pathways were detected by searching with K numbers in the KEGG pathway database.

### 2.5. Prediction of subcellular localization

Prediction of the subcellular location of proteins is a significant step in finding potential drug targets. PSORTb v3.0 was used to foretell whether a protein is located in the cytoplasm, cytoplasmic membrane, cell wall, or extracellular space based on localization scores. PSORTb 3.0 is claimed to be the most precise SCL predictor that has increased recall and predictive coverage. It uses a cutoff of 7.5 as a reliable threshold for assigning a single localization (Yu et al., 2010).

### 2.6. Druggability analysis

Human non-homologous essential cytoplasmic clusters that are involved in the unique metabolic pathways of *Acinetobacter* were subjected to BLASTp against all drug target sequences available at DrugBank Database with a threshold of E value 10^−5^ (Wishart et al., 2006; Ashraf et al., 2022). The proteins with ‘Hits’ and ‘No hits’ were considered as druggable and novel targets, respectively.

### 2.7. Primary and secondary structure analysis

Primary structures of novel target protein clusters were evaluated using the Expasy ProtParam tool (Gasteiger et al., 2003). ProtParam tool computes various physiochemical parameters including molecular weight, theoretical pI, instability index, aliphatic index, and grand average of hydropathicity (GRAVY). Secondary structures of target proteins were predicted by SOPMA (Geourjon and Deleage, 1995). SOPMA provides various information on the different conformational states of protein including α-helices, β-turns, extended loops, and random coils.

### 2.8. 3D structure prediction and validation

SWISS-MODEL was used to perform homology modeling to predict the three-dimensional structure of novel target proteins. GMQE (Global Model Quality Estimate) is a measurement of the model quality based on the template structure and target-template alignment. Templates from AlphaFold DB (usually exhibit high quality suitable for homology modeling) were considered to build models due to better coverage and higher GMQE score than experimentally determined templates (Schwede et al., 2003). These tertiary structures were validated by the SAVES server on the basis of PROCHECK. PROCHECK assesses the stereochemical quality of 3D structures of protein (Laskowski et al., 1993).

### 2.9. Evaluation of druggability of the novel targets

DoGSiteScorer was employed to estimate the druggability of novel target proteins. PDB files of 3D models were subjected to DoGSiteScorer. This server detects and estimates the druggability of protein pockets. Ligands are expected to bind with high affinity to pockets with a higher druggability score. It also computes volume, surface area, depth, and some other metrics for each predicted pocket (Volkamer et al., 2012).

### 2.10. Structural alignment and comparison

MatchMaker of UCSF Chimera 1.18 was used to compare proteins by creating their superimposed structures (Meng et al., 2006). MatchMaker performs structural superimposition by starting with pairwise sequence alignments and subsequently fitting the aligned residue pairs. Additionally, Match -> Align tool from structure comparison was employed to generate a sequence alignment from the structural superposition of proteins (Meng et al., 2006). Information regarding the percent identities of the proteins were obtained from the alignment. Furthermore, amino acid residues of the common drug-binding pockets were closely observed both in the superimposed structures and the sequence alignment.

## 3. Results & Discussion

### 3.1. Core-genes from bacteria

In order to target multiple pathogenic species, it is essential to target specific proteins that are common to all those species. Pan-genome comprises of core-genome, dispensable genome, and species-specific genes. Core-genome contains genes that are present in all genomes of a particular dataset. The majority of these genes play vital roles in the survival of bacteria (Costa et al., 2020; Inglin et al., 2018). A study even showed that the core genes are crucial to *Acinetobacter baumannii*’s ability to resist tobramycin (Gallagher et al., 2017). Pan-genome Explorer returned 1245 core-gene clusters as output, where every cluster had genes from each of the four *Acinetobacter* species. The cluster counts at each step are outlined in Table 1 and illustrated graphically in Fig. 2. Only core protein sequences were considered for subsequent analysis.

**Fig. 2.**
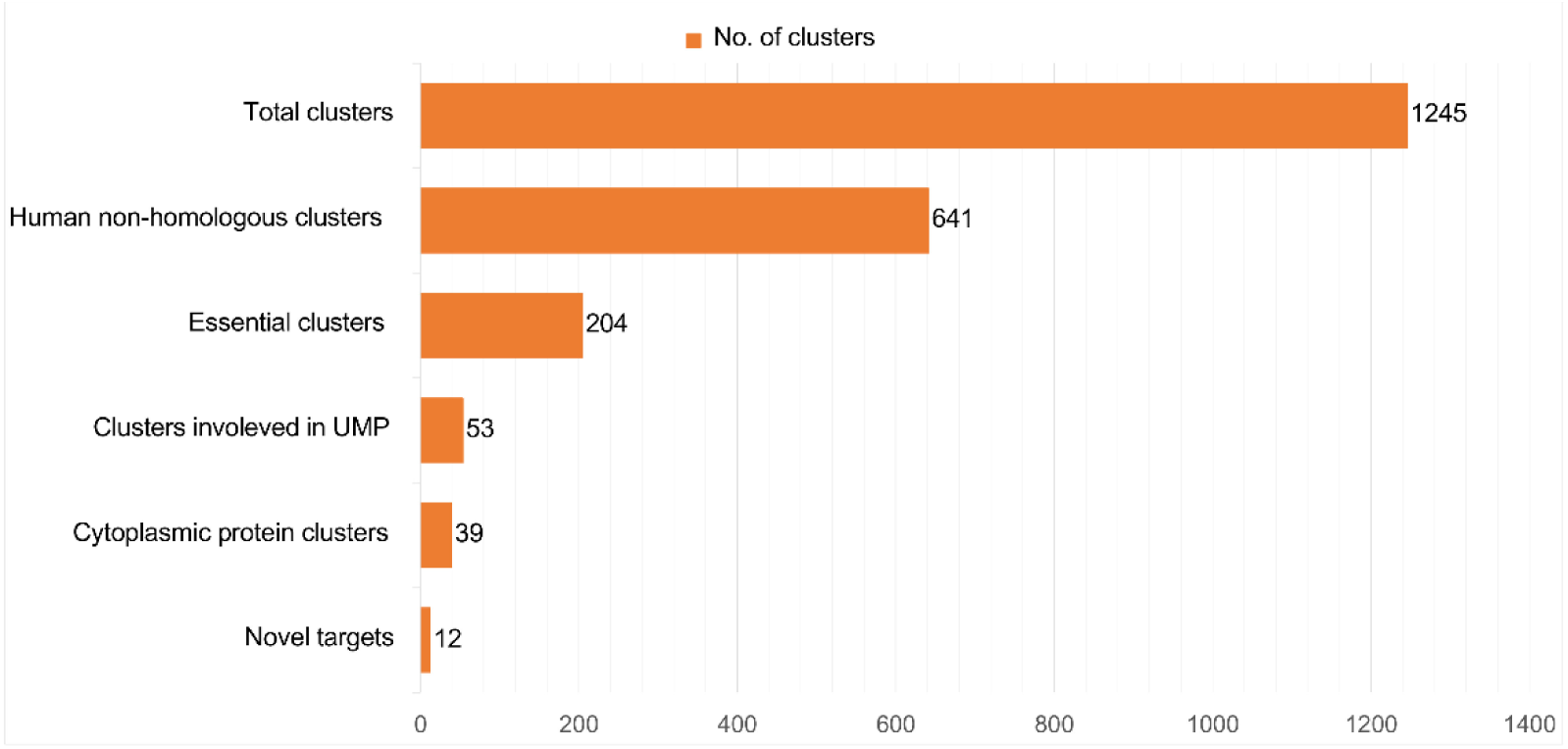
Protein cluster counts at each step. The orange bars represent the number of clusters.

**Table 1.** Analytical steps of subtractive genome analysis.

| Analytical Steps | Number of Core-gene Clusters |
| --- | --- |
| Clusters collected from Panexplorer | 1245 |
| Human non-homologous clusters (E-value $10^{-3}$) | 641 |
| Essential clusters in DEG (E value $\leq 10^{-100}$, Bit score $> 100$) | 204 |
| Essential clusters participate in metabolic pathways (KEGG) | 134 |
| Essential clusters participate in unique metabolic pathways (KEGG) | 53 |
| Human non-homologous essential cytoplasmic protein clusters | 39 |
| Clusters that matched with drug target sequences of DrugBank (E-value $10^{-5}$) | 26 |
| Novel drug target clusters | 12 |

### 3.2. Human non-homologous clusters

During drug development, we need to ensure that therapeutics do not cross-react with homologous proteins in humans. Using homologous proteins as drug targets during treatment may cause host cytotoxicity (Hossain et al., 2017). Through a similarity search of 1245 core-gene clusters against human RefSeq proteins (GRCh38) using BLASTp, 641 clusters were found to be non-homologous to human proteome. Since these clusters were unique in pathogens (*Acinetobacter* spp.), they were potential drug targets.

### 3.3. Essential clusters in DEG

Essential genes are vital for bacteria to survive in particular conditions (Butt et al., 2012). Disrupting the functions of essential proteins will result in cell death. Therefore, targeting these essential proteins can enhance the therapeutic efficacy of treating bacterial infections (Yan and Gao, 2020). After BLASTp analysis of human non-homologous clusters against essential bacterial proteins of DEG, 204 out of 641 clusters were identified to be essential for the survival of *Acinetobacter*. The rest of the clusters were omitted from the study since targeting these clusters may not be effective in killing or inhibiting the activity of selected *Acinetobacter* species.

### 3.4. Clusters involved in common and unique metabolic pathways

Unique pathways refer to the pathways that are present in pathogen but are missing in the host. Proteins involved in these pathways could be useful as potential drug and vaccine targets (Butt et al., 2012). 134 core-gene clusters participated in different metabolic pathways according to KEGG analysis. Among them, 53 clusters were involved in unique pathways. The rest of the clusters were found in common metabolic pathways. It is risky to target proteins that participate in common metabolic pathways since such pathways are found in both pathogen and human, which may lead to potential side effects (Oany et al., 2018). Therefore, 53 clusters (212 protein sequences) that were solely involved in unique pathways were considered for further investigation (Table 2).

**Table 2.** Protein clusters involved in unique metabolic pathways.

| Serial | Unique Pathway Name | Pathway ID | KEGG Orthology |
| --- | --- | --- | --- |
| 1 | Bacterial secretion system | 03070 | K02454 K02455 K03070 K03072<br>K03074 K03076 |
| 2 | Beta-Lactam resistance | 01501 | K03585 K03587 K18139 |
| 3 | Biosynthesis of various plant secondary metabolites | 00999 | K00014 |
| 4 | C5-Branched dibasic acid metabolism | 00660 | K01653 |
| 5 | Cationic antimicrobial peptide (CAMP) resistance | 01503 | K03585 |
| 6 | Flagellar assembly | 02040 | K03086 K03092 |
| 7 | Lipopolysaccharide biosynthesis | 00540 | K00748 K00979 K01627 K02535 |
| 8 | Lysine biosynthesis | 00300 | K00003 K00133 K00215 K00674<br>K00928 K01778 K01928 K01929 |
| 9 | Methane metabolism | 00680 | K00925 K01007 K01595 K01624<br>K15633 |
| 10 | Monobactam biosynthesis | 00261 | K00133 K00215 K00928 K00957 |
| 11 | Novobiocin biosynthesis | 00401 | K24018 |
| 12 | O-Antigen nucleotide sugar biosynthesis | 00541 | K01840 |
| 13 | Peptidoglycan biosynthesis | 00550 | K00790 K01000 K01924 K01925<br>K01928 K01929 K02563 K03587<br>K07258 |
| 14 | Quorum sensing | 02024 | K01497 K01626 K01657 K03070<br>K03076 K18139 |
| 15 | Two-component system | 02020 | K00425 K00426 K00990 K02313<br>K03092 K07639 K07659 K07661<br>K07678 K07689 |
| 16 | Vancomycin resistance | 01502 | K01000 K01929 K02563 |

### 3.5. Identification of cytoplasmic protein clusters

The localization of a protein within a cell reveals information about its location and function (Fatoba et al., 2021). After the analysis of the remaining clusters in pSORTb server, 39 out of 53 clusters were found to be cytoplasmic proteins. The rest of the clusters were cytoplasmic membrane and outer membrane proteins. It is difficult to purify and assay proteins that are localized in membranes (Duffield et al., 2010). Therefore, cytoplasmic proteins are usually a good choice as drug targets whereas membrane proteins are utilized for vaccine production (Ashraf et al., 2022). Moreover, cytoplasmic proteins are easily accessible to drugs (Khan et al., 2021). Only the cytoplasmic protein clusters were taken to the next step of analysis.

### 3.6. Identification of novel targets

Due to resistance to existing drugs, finding new targets is now a priority (Gupta et al., 2019). The BLASTp result of 39 cytoplasmic clusters against drug target proteins of DrugBank revealed the similarity of 26 clusters. Rest of the clusters (12) did not show significant similarity based on the specific threshold. These 12 clusters that did not match with the drug target proteins of the DrugBank database were considered novel targets. The KEGG Orthology numbers and corresponding pathways of these clusters are represented in Table 3. Additionally, Table 4 presents the roles of the proteins from novel target clusters. 26 clusters that matched with the database were druggable targets. In a particular cluster, four of the sequences were neither novel nor druggable. Since the BLASTp result differed among these sequences, that specific cluster was excluded from further analysis.

**Table 3.** Novel drug target clusters involved in different unique metabolic pathways.

| KEGG Orthology | Unique Metabolic Pathway |
| --- | --- |
| K00003 K01778 K00928 | Lysine biosynthesis |
| K00928 K00957 | Monobactam biosynthesis |
| K00990 K02313 K03092 | Two-component system |
| K03092 | Flagellar assembly |
| K00748 | Lipopolysaccharide biosynthesis |
| K00925 | Methane metabolism |
| K01497 K03070 | Quorum sensing |
| K01653 | C5-Branched dibasic acid metabolism |
| K03070 | Bacterial secretion system |

**Table 4.** Functions of novel protein clusters.

| Cluster Serial No. | Protein Name* | Role |
| --- | --- | --- |
| 1 | Homoserine dehydrogenase | Regulates aspartate pathway; facilitates the synthesis of lysine, methionine, threonine, and isoleucine (Kim et al., 2020) |
| 2 | Diaminopimelate (DAP) epimerase | Participates in the biosynthesis of meso-DAP and lysine (precursors for peptidoglycan, housekeeping protein, and virulence factor synthesis) (Hor et al., 2013) |
| 3 | Aspartate kinase | Participates in the biosynthesis of aspartate-derived amino acids (Min et al., 2015) |
| 4 | Sulfate adenylyltransferase | Activates inorganic sulfate and incorporate it into organic compounds (Ullrich et al., 2001) |
| 5 | [protein-PII] uridylyltransferase | Uridylylates and de-uridylylates the small trimeric protein PII (Bairoch, 2000) |
| 6 | Replication initiator protein | Initiates the DNA replication process by recognizing and binding to the replicator (Stenlund, 2003) |
| 7 | RNA polymerase factor sigma-54 | Binds to RNA polymerase and regulates the transcription of various functional genes (Yu et al., 2021) |
| 8 | Lipid A-disaccharide synthase | Participates in the biosynthesis of the phosphorylated glycolipid, lipid A, in the outer membrane of Gram negative bacteria (Bairoch, 2000) |
| 9 | Acetate kinase | Essential in anaerobic metabolism, catalyzes the reversible phosphorylation of acetate by ATP (Ingram-Smith et al., 2015) |
| 10 | GTP cyclohydrolase II | Participates in the biosynthesis of riboflavin (Ritz et al., 2001) |
| 11 | Acetolactate synthase | Participates in the biosynthesis of leucine, isoleucine, and valine (Smith et al., 1989) |
| 12 | Preprotein translocase | Facilitates the movement of secretory proteins across membranes (Economou, 1998) |

\* Protein names have been generalized to denote primary functional roles.

### 3.7. Primary and secondary structures of novel clusters

Instability index, molecular weight, theoretical pI, and grand average hydropathicity (GRAVY) scores were considered to determine potential drug target clusters. A score of instability index equal to or above 40 indicates that the particular protein is unstable. Positive GRAVY score means the nature of a protein is hydrophobic whereas negative GRAVY score means the protein is hydrophilic. The cutoff values followed in this study were: instability index < 40; molecular weight < 100K Da; GRAVY > -0.240; theoretical pI < 7.2 (Hafsa et al., 2022). Among 12 protein clusters, 5 clusters (cluster 1, 2, 3, 9, and 11) were selected and 7 clusters were rejected (provided in Table 5). The percentages of secondary structures for the selected five clusters have been provided in Table 6.

**Table 5.**
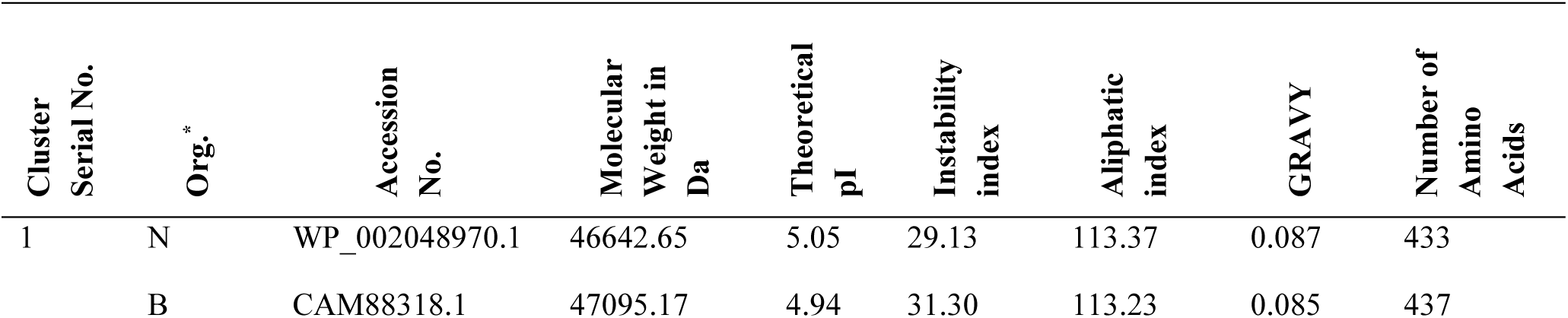
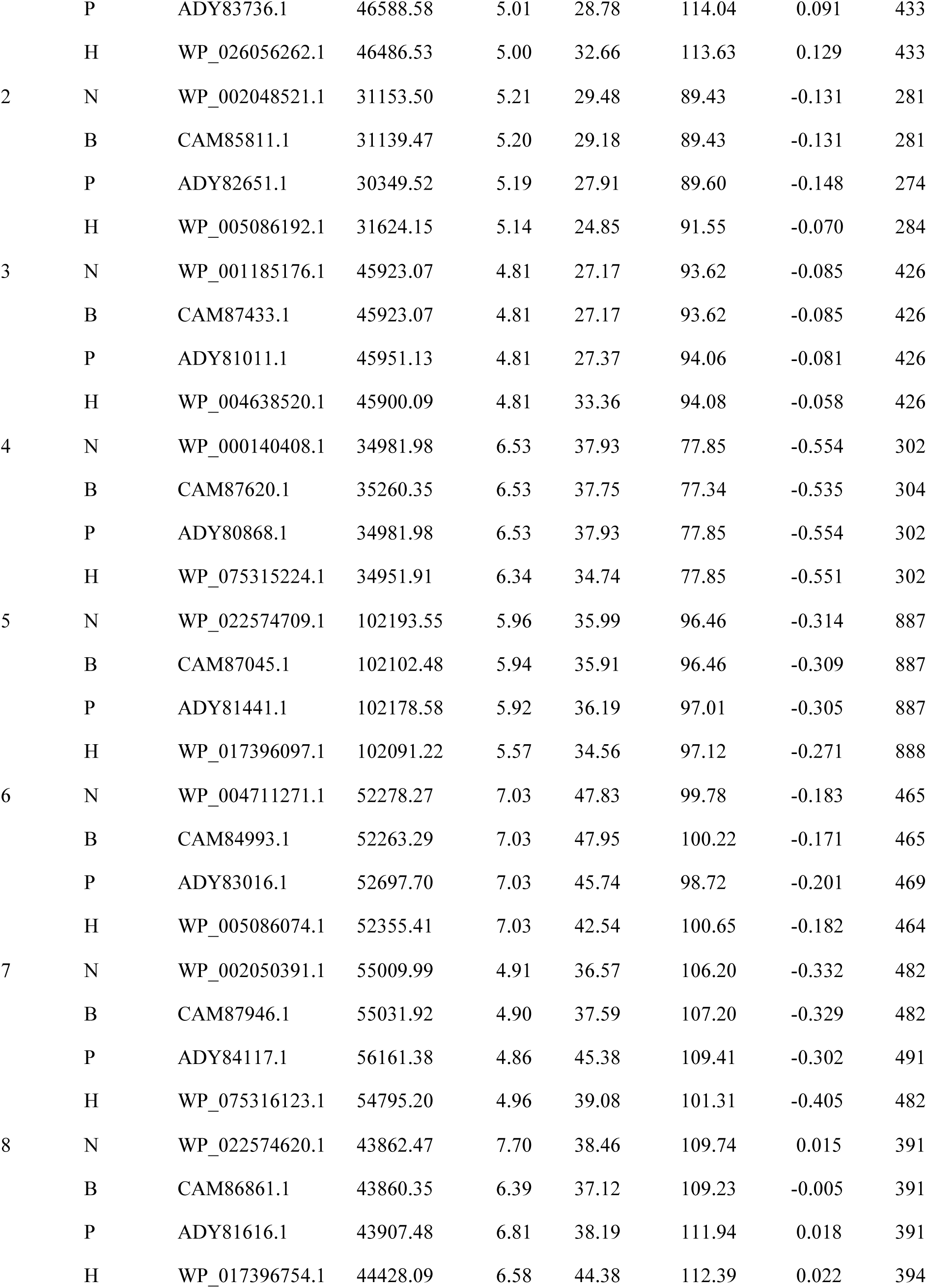
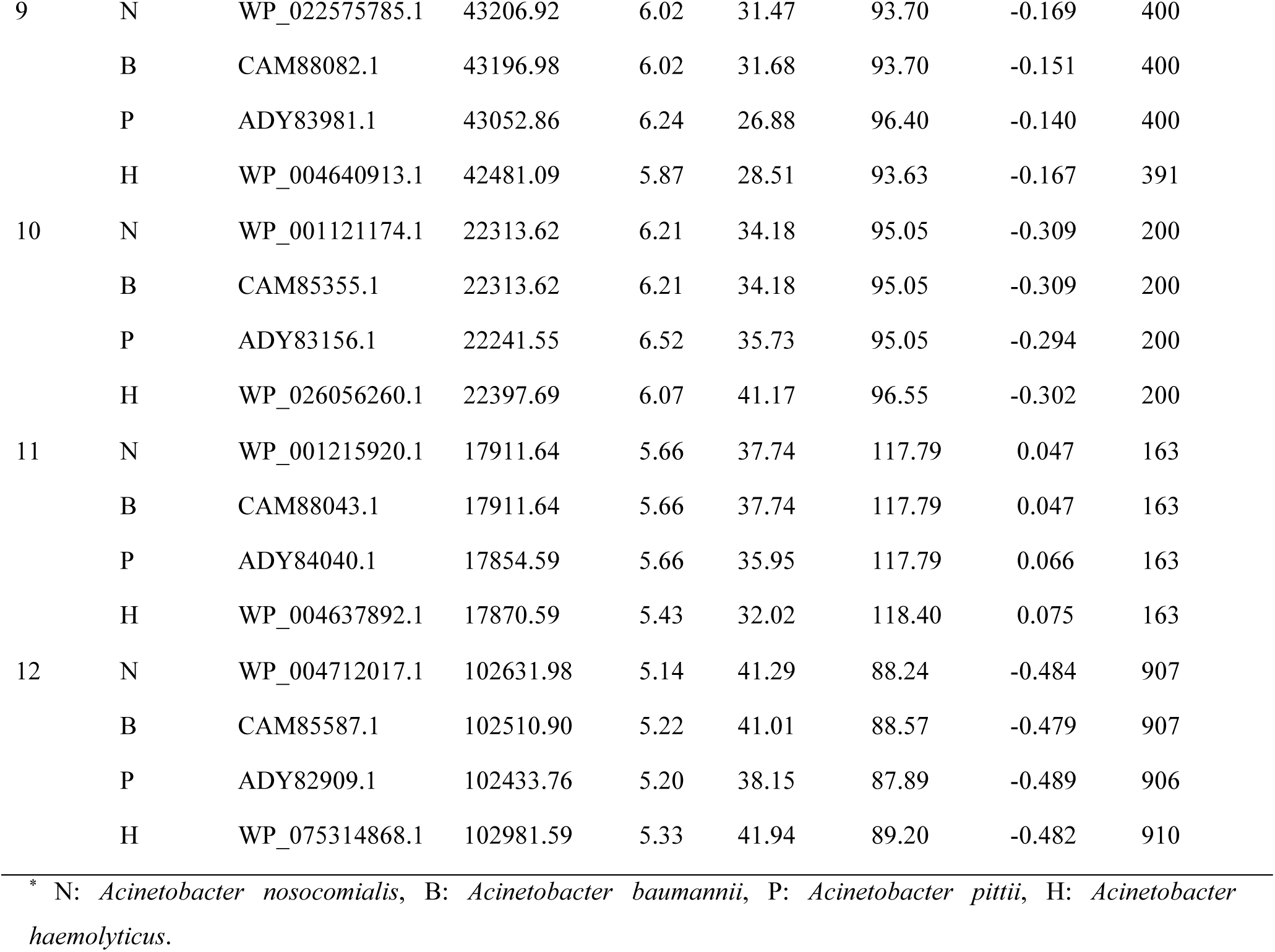
Information about the primary structures of novel target clusters.

**Table 6.** Secondary structure prediction of novel drug target clusters.

| Cluster Serial | Accession No. | Alpha Helix | Beta Turns | Extended Strand | Random Coils |
| --- | --- | --- | --- | --- | --- |
| 1 | WP_002048970.1 | 41.80% | 8.55% | 17.55% | 32.10% |
|  | CAM88318.1 | 43.71% | 8.01% | 17.62% | 30.66% |
|  | ADY83736.1 | 43.65% | 7.39% | 17.55% | 31.41% |
|  | WP_026056262.1 | 44.11% | 8.08% | 16.86% | 30.95% |
| 2 | WP_002048521.1 | 13.52% | 9.61% | 33.10% | 43.77% |
|  | CAM85811.1 | 14.59% | 9.61% | 33.81% | 41.99% |
|  | ADY82651.1 | 18.25% | 10.58% | 31.39% | 39.78% |
|  | WP_005086192.1 | 18.66% | 8.45% | 30.63% | 42.25% |
| 3 | WP_001185176.1 | 37.79% | 7.75% | 21.83% | 32.63% |
|  | CAM87433.1 | 37.79% | 7.75% | 21.83% | 32.63% |
|  | ADY81011.1 | 36.85% | 6.34% | 22.54% | 34.27% |
|  | WP_004638520.1 | 38.97% | 7.28% | 21.83% | 31.92% |
| 9 | WP_022575785.1 | 37.75% | 7.75% | 21.00% | 33.50% |
|  | CAM88082.1 | 38.00% | 9.25% | 19.75% | 33.00% |
|  | ADY83981.1 | 40.75% | 8.75% | 18.25% | 32.25% |
|  | WP_004640913.1 | 39.13% | 9.21% | 19.69% | 31.97% |
| 11 | WP_001215920.1 | 41.10% | 7.98% | 27.61% | 23.31% |
|  | CAM88043.1 | 41.10% | 7.98% | 27.61% | 23.31% |
|  | ADY84040.1 | 44.79% | 5.52% | 26.38% | 23.31% |
|  | WP_004637892.1 | 41.72% | 6.75% | 26.99% | 24.54% |

### 3.8. Homology modeling

Three-dimensional structures of the five selected protein clusters were predicted by homology modeling in the SWISS-MODEL server. The predicted structures were verified by PROCHECK in the SAVES server. More than 90% of residues are expected to be in the most favored regions in case of good quality models. Above 90% of the residues of the four proteins of cluster 1 (i.e. WP_002048970.1, CAM88318.1, ADY83736.1, WP_026056262.1) were in the most favoured regions in Ramachandran plot (Fig. 3). For WP_002048970.1, CAM88318.1, and ADY83736.1, only a single residue fell in the disallowed region, whereas WP_026056262.1 had two residues in that region.

**Fig. 3.**
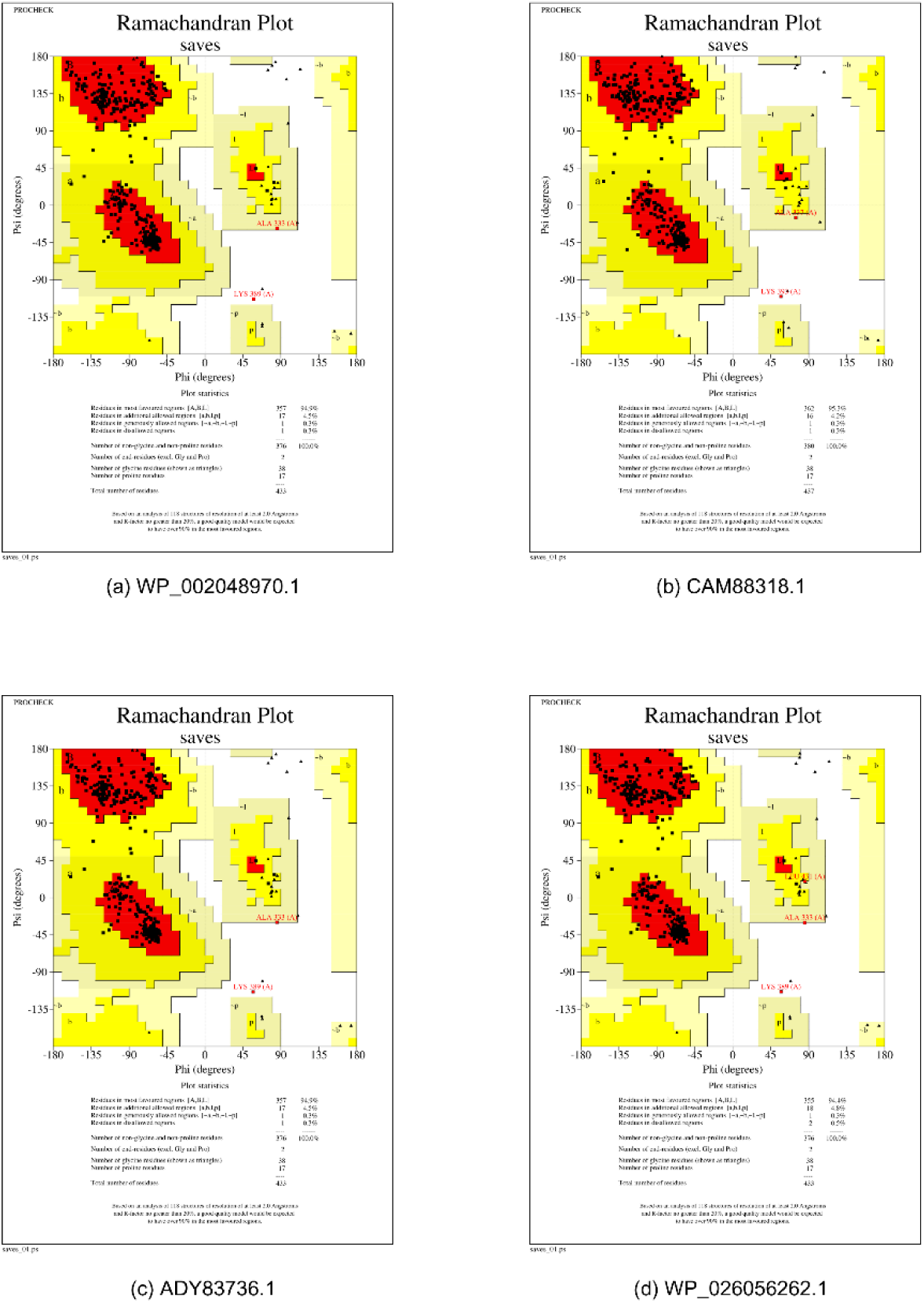
Ramachandran plots of the four proteins in cluster 1. (a) WP_002048970.1 from *Acinetobacter nosocomialis*, (b) CAM88318.1 from *Acinetobacter baumannii*, (c) ADY83736.1 from *Acinetobacter pittii*, and (d) WP_026056262.1 from *Acinetobacter haemolyticus*.

### 3.9. Druggability analysis and Structural Comparison

Among the five clusters, each protein of cluster 1 had a drug binding pocket located in nearly the same position (Fig. 4). During druggability analysis, pockets that have druggability score ≥ 0.8 were considered to have better druggability (Hassan et al., 2023). The proteins of cluster 1 had high druggability scores, ranging from 0.80 to 0.87. DoGSiteScorer calculation results have been presented in Table 7. The superimposed structures of them showed that they are nearly identical. Binding sites of the common drug-binding pockets have been shown in Fig. 5 and in ‘Supplementary File S1’. Sequence alignment from the structural superposition of these proteins revealed that WP_002048970.1 was 99.08%, 99.08%, and 96.07% identical to CAM88318, ADY83736.1, and WP_026056262.1 respectively (Fig. 6).

**Fig. 4.**
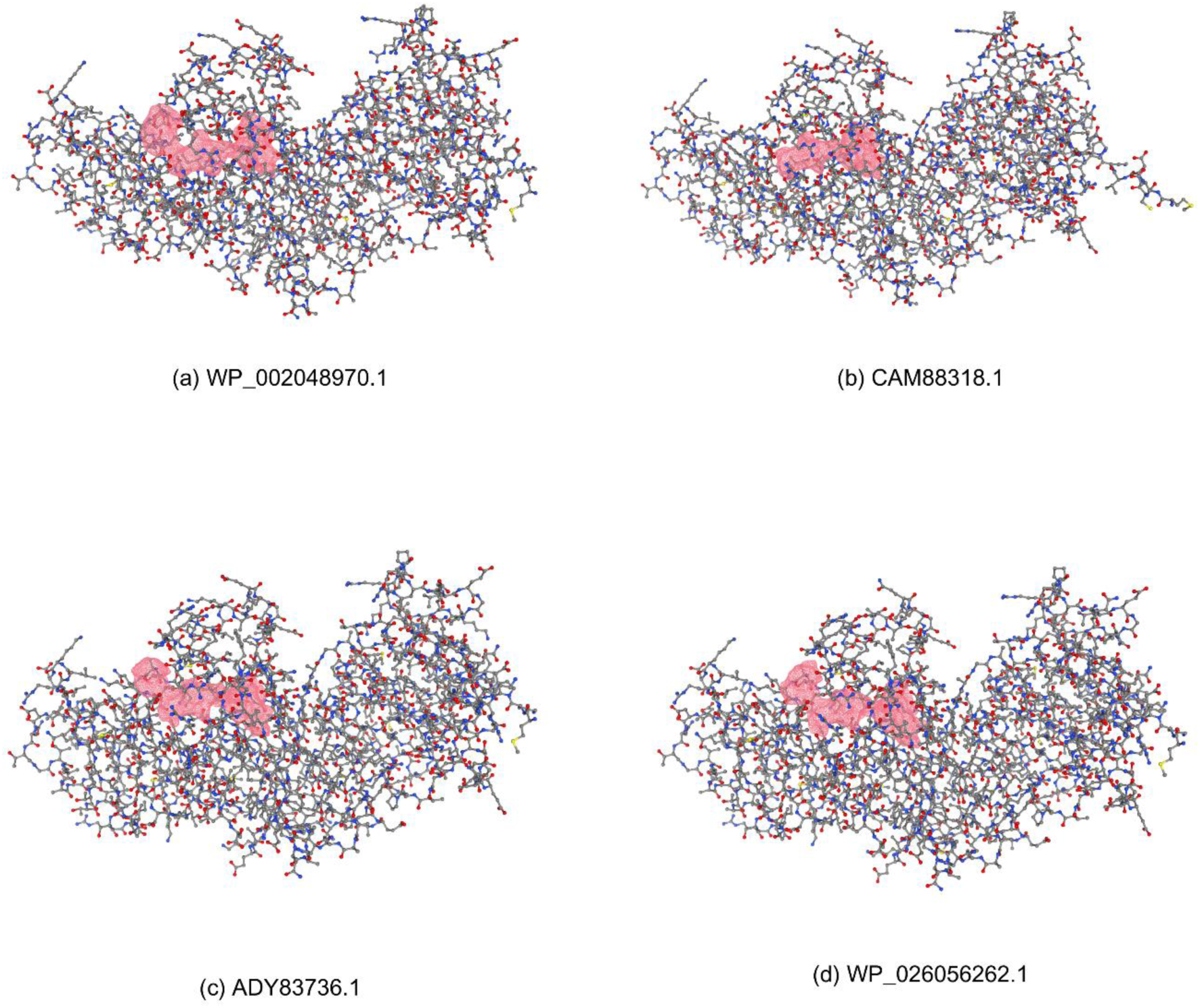
Ball-and-stick models of the four proteins in cluster 1. (a) WP_002048970.1 from *Acinetobacter nosocomialis*, (b) CAM88318.1 from *Acinetobacter baumannii*, (c) ADY83736.1 from *Acinetobacter pittii*, and (d) WP_026056262.1 from *Acinetobacter haemolyticus*. Potential common drug-binding pockets are shown in red.

**Fig. 5.**
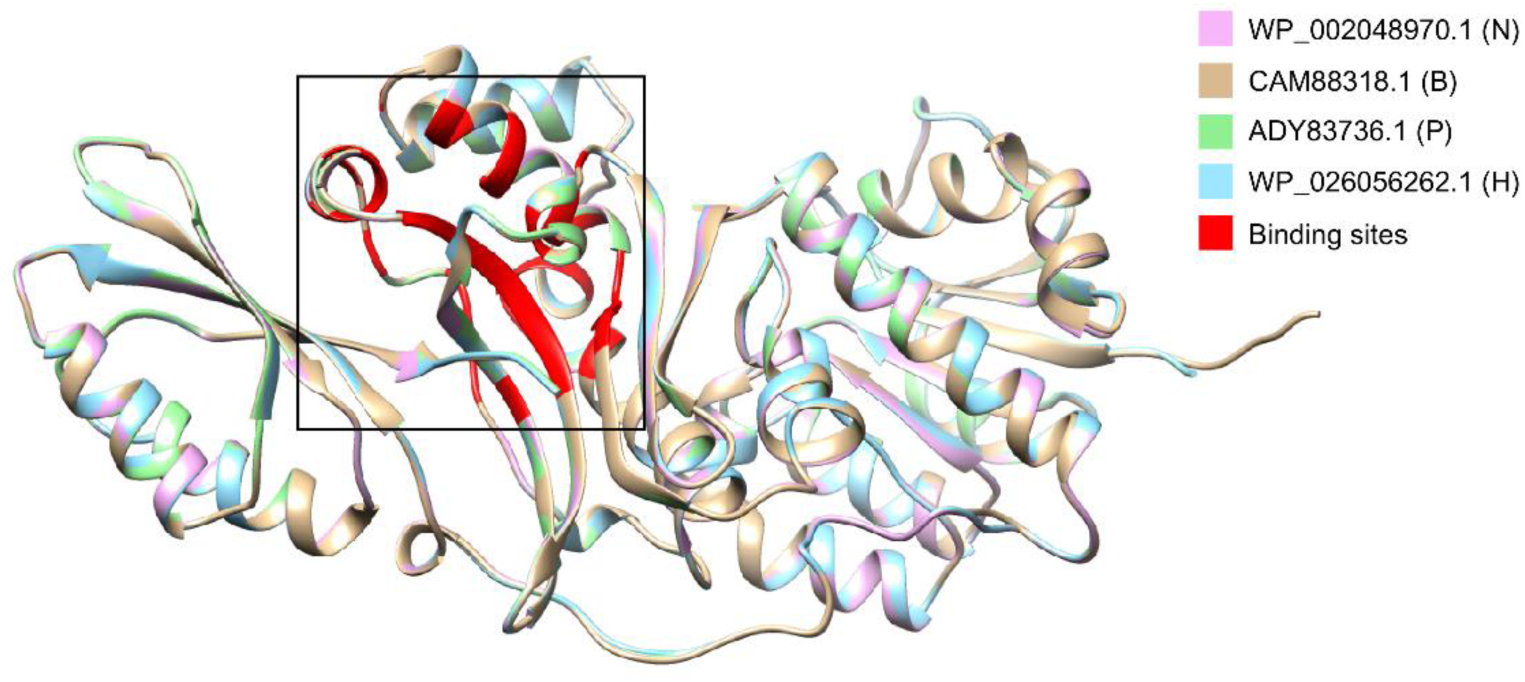
Structural superimposition of cluster 1 proteins using Chimera. Binding sites of the common drug-binding pockets are shown in the box (red).

**Fig. 6.**
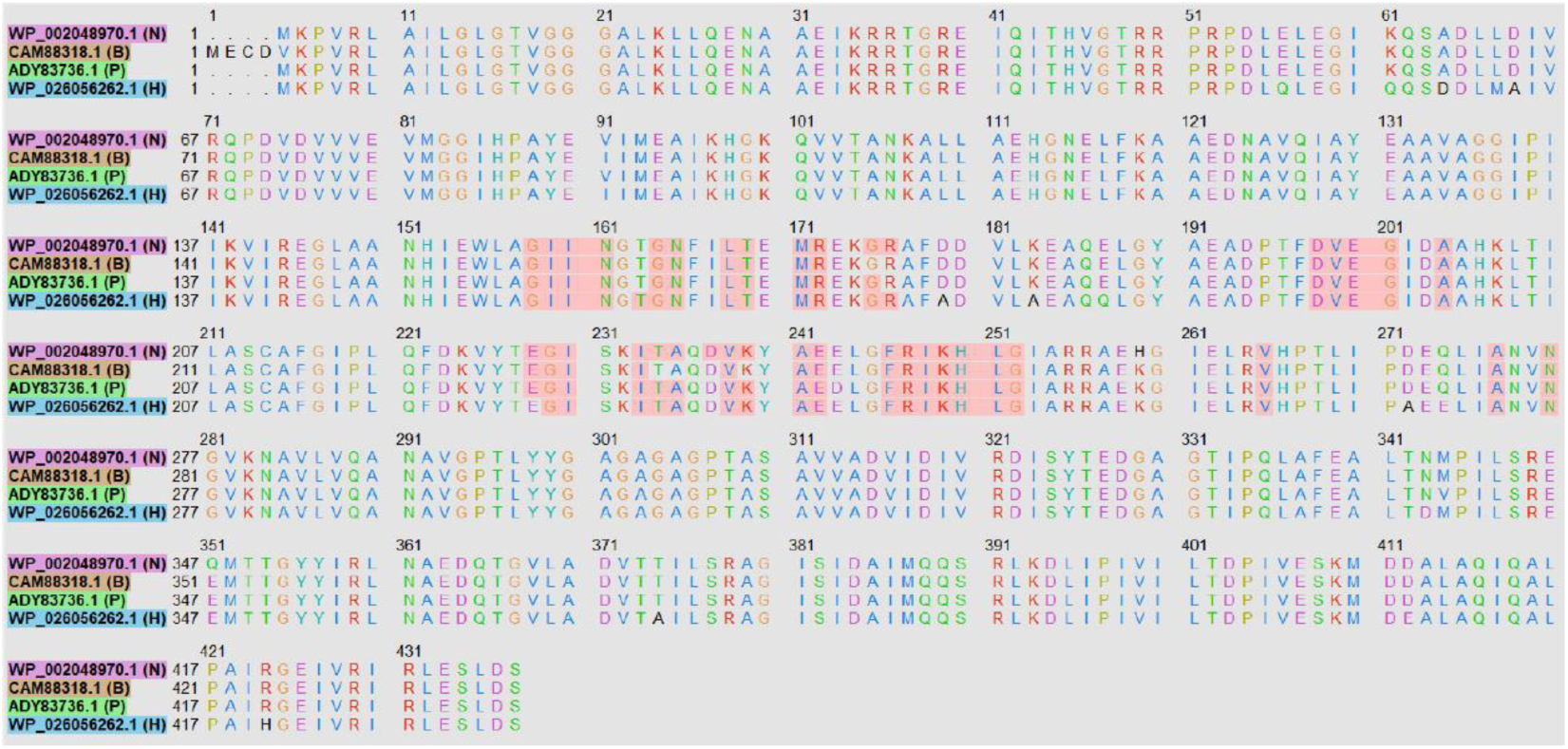
Sequence alignment from the structural superposition of cluster 1 proteins. Binding residues of the common drug-binding pockets are shaded in red.

**Table 7.** DoGSiteScorer calculation results for proteins in cluster 1.

| Accession ID (Cluster 1) | Pocket Name | Volume Å<sup>3</sup> | Surface Å<sup>2</sup> | Drug Score |
| --- | --- | --- | --- | --- |
| WP_002048970.1 | P_3 | 419.54 | 463.72 | 0.85 |
| CAM88318.1 | P_3 | 321.42 | 306.79 | 0.80 |
| ADY83736.1 | P_3 | 400.66 | 433.79 | 0.85 |
| WP_026056262.1 | P_3 | 406.72 | 435.88 | 0.87 |

The proteins of cluster 1 were homoserine dehydrogenases. It is an essential regulatory protein that participates in the biosynthesis of amino acids (Ogata et al., 2018). Homoserine dehydrogenase (KEGG Entry: K00003) is involved in lysine biosynthesis which is a unique pathway present in the bacteria and absent in human. Humans obtain lysine from diet (Schobel et al., 2010). Lysine/DAP biosynthetic pathway is an attractive target because of its significant role in biosynthesis of cell wall and amino acid (Garg et al., 2010). Homoserine dehydrogenase can be inhibited to disrupt this pathway. Moreover, it is a promising pathway for novel antibiotic targets (Muduli et al., 2023).

## 4. Conclusion

Subtractive genomics is a widely utilized method to subtract protein sequences from pathogen proteomes on the basis of several criteria such as host non-homogeneity, essentiality, specific subcellular localization, participation in unique metabolic pathways, and other considerations. This approach enables researchers to identify drug targets from large datasets of proteins. The present study identified four proteins (WP_002048970.1, CAM88318.1, ADY83736.1, WP_026056262.1) that have high druggability and similar drug target regions. Inhibition of these cytoplasmic proteins (homoserine dehydrogenases) might help in managing *Acinetobacter* infections. Furthermore, this cluster-based method of subtractive genomics can also be utilized to find potential drug targets in other organisms.

## Supporting information

S1. Common drug-binding pockets (red) of cluster 1 proteins.

## Supplementary material

**S1.** Common drug-binding pockets (red) of cluster 1 proteins.

## CRediT authorship contribution statement

**Soaibur Rahman:** Conceptualization, Data curation, Formal analysis, Methodology, Writing – original draft, Writing – review and editing. **Soharth Hasnat:** Conceptualization, Methodology, Writing – review & editing. **M Murshida Mahbub:** Conceptualization, Methodology, Writing – review & editing. **Zinat Farzana:** Conceptualization. **Sal Sabila:** Formal analysis.

## Acknowledgments

We are thankful to Shahriar Mahmud from East West University for providing technical resources during the last phase of this study.

## References

[1] S.B. Almasaudi, *Acinetobacter* spp. as nosocomial pathogens: Epidemiology and resistance features, Saudi J Biol Sci. 25 (2018) 586–596. 10.1016/j.sjbs.2016.02.009.

[2] H. Wisplinghoff, M.B. Edmond, M.A. Pfaller, R.N. Jones, R.P. Wenzel, H. Seifert, Nosocomial bloodstream infections caused by *Acinetobacter* species in United States hospitals: clinical features, molecular epidemiology, and antimicrobial susceptibility, Clin Infect Dis. 31 (2000) 690–697. 10.1086/314040.

[3] E. Bergogne-Bérézin, Importance of Acinetobacter spp, in: E. Bergogne-Bérézin, H. Friedman, M. Bendinelli (Eds.), Acinetobacter Biology and Pathogenesis, Springer US, New York, NY, 2008, pp. 1–18.

[4] J. Kiskova, A. Juhas, S. Galuskova, L. Malinicova, M. Kolesarova, M. Piknova, P. Pristas, Antibiotic Resistance and Genetic Variability of *Acinetobacter* spp. from Wastewater Treatment Plant in Koksov-Baksa (Kosice, Slovakia), Microorganisms. 11 (2023). 10.3390/microorganisms11040840.

[5] Centers for Disease Control and Prevention, About Acinetobacter. https://www.cdc.gov/Acinetobacter/about/index.html, 2024 (accessed 3 September 2024).

[6] A. Lupo, M. Haenni, J.Y. Madec, Antimicrobial Resistance in *Acinetobacter* spp. and *Pseudomonas* spp, Microbiol Spectr. 6 (2018). 10.1128/microbiolspec.ARBA-0007-2017.

[7] I. Cavallo, A. Oliva, R. Pages, F. Sivori, M. Truglio, G. Fabrizio, M. Pasqua, F. Pimpinelli, E.G. Di Domenico, *Acinetobacter baumannii* in the critically ill: complex infections get complicated, Front Microbiol. 14 (2023) 1196774. 10.3389/fmicb.2023.1196774.

[8] F. Perez, A.M. Hujer, K.M. Hujer, B.K. Decker, P.N. Rather, R.A. Bonomo, Global challenge of multidrug-resistant *Acinetobacter baumannii*, Antimicrob Agents Chemother. 51 (2007) 3471–3484. 10.1128/AAC.01464-06.

[9] A. Nithichanon, C. Kewcharoenwong, H. Da-Oh, S. Surajinda, A. Khongmee, S. Koosakunwat, B.W. Wren, R.A. Stabler, J.S. Brown, G. Lertmemongkolchai, *Acinetobacter nosocomialis* Causes as Severe Disease as *Acinetobacter baumannii* in Northeast Thailand: Underestimated Role of *A. nosocomialis* in Infection, Microbiol Spectr. 10 (2022) e0283622. 10.1128/spectrum.02836-22.

[10] H. Pailhories, C. Tiry, M. Eveillard, M. Kempf, *Acinetobacter pittii* isolated more frequently than *Acinetobacter baumannii* in blood cultures: the experience of a French hospital, J Hosp Infect. 99 (2018) 360–363. 10.1016/j.jhin.2018.03.019.

[11] B.T. Tierney, N.K. Singh, A.C. Simpson, A.M. Hujer, R.A. Bonomo, C.E. Mason, K. Venkateswaran, Multidrug-resistant *Acinetobacter pittii* is adapting to and exhibiting potential succession aboard the International Space Station, Microbiome. 10 (2022) 210. 10.1186/s40168-022-01358-0.

[12] L. Bai, S. Zhang, Y. Deng, C. Song, G. Kang, Y. Dong, Y. Wang, F. Gao, H. Huang, Comparative genomics analysis of *Acinetobacter haemolyticus* isolates from sputum samples of respiratory patients, Genomics. 112 (2020) 2784–2793. 10.1016/j.ygeno.2020.03.016.

[13] M.A. Islam, M.S. Hossain, S. Hasnat, M.H. Shuvo, S. Akter, M.A. Maria, A. Tahcin, M.A. Hossain, M.N. Hoque, Scientific Reports, 14 (2024) 17182. 10.1038/s41598-024-65112-2

[14] M.S. Hossain, S. Hasnat, S. Akter, M.M. Mim, A. Tahcin, M. Hoque, D. Sutradhar, M.A.A. Keya, N.R. Sium, S. Hossain, R. Masuma, S.H. Rakib, M.A. Islam, T. Islam, P. Bhattacharya, M.N. Hoque, Front Pharmacol, 15 (2024) 1465827. 10.3389/fphar.2024.1465827

[15] S. Hasnat, M.N. Hoque, M.M. Mahbub, T.I. Sakif, A.D.A. Shahinuzzaman, T. Islam, Pantothenate kinase: A promising therapeutic target against pathogenic *Clostridium* species, Heliyon. 10 (2024) e34544. 10.1016/j.heliyon.2024.e34544.

[16] C. UniProt, UniProt: the universal protein knowledgebase in 2021, Nucleic Acids Res. 49 (2021) D480–D489. 10.1093/nar/gkaa1100.

[17] A. Dereeper, M. Summo, D.F. Meyer, PanExplorer: a web-based tool for exploratory analysis and visualization of bacterial pan-genomes, Bioinformatics. 38 (2022) 4412–4414. 10.1093/bioinformatics/btac504.

[18] A. Wadood, A. Jamal, M. Riaz, A. Khan, R. Uddin, M. Jelani, S.S. Azam, Subtractive genome analysis for *In silico* identification and characterization of novel drug targets in *Streptococcus pneumonia* strain JJA, Microb Pathog. 115 (2018) 194–198. 10.1016/j.micpath.2017.12.063.

[19] S.F. Altschul, W. Gish, W. Miller, E.W. Myers, D.J. Lipman, Basic local alignment search tool, J Mol Biol. 215 (1990) 403–410. 10.1016/S0022-2836(05)80360-2.

[20] M. Sarkar, L. Maganti, N. Ghoshal, C. Dutta, *In silico* quest for putative drug targets in *Helicobacter pylori* HPAG1: molecular modeling of candidate enzymes from lipopolysaccharide biosynthesis pathway, J Mol Model. 18 (2012) 1855–1866. 10.1007/s00894-011-1204-3.

[21] A. Rahman, S. Noore, A. Hasan, R. Ullah, H. Rahman, A. Hossain, Y. Ali, S. Islam, Identification of potential drug targets by subtractive genome analysis of *Bacillus anthracis* A0248: An *In silico* approach, Comput Biol Chem. 52 (2014) 66–72. 10.1016/j.compbiolchem.2014.09.005.

[22] Y.T. Liang, H. Luo, Y. Lin, F. Gao, Recent advances in the characterization of essential genes and development of a database of essential genes, Imeta. 3 (2024) e157. 10.1002/imt2.157.

[23] R. Zhang, H.Y. Ou, C.T. Zhang, DEG: a database of essential genes, Nucleic Acids Res. 32 (2004) D271–272. 10.1093/nar/gkh024.

[24] M. Kanehisa, S. Goto, KEGG: kyoto encyclopedia of genes and genomes, Nucleic Acids Res. 28 (2000) 27–30. 10.1093/nar/28.1.27.

[25] N.Y. Yu, J.R. Wagner, M.R. Laird, G. Melli, S. Rey, R. Lo, P. Dao, S.C. Sahinalp, M. Ester, L.J. Foster, F.S. Brinkman, PSORTb 3.0: improved protein subcellular localization prediction with refined localization subcategories and predictive capabilities for all prokaryotes, Bioinformatics. 26 (2010) 1608–1615. 10.1093/bioinformatics/btq249.

[26] D.S. Wishart, C. Knox, A.C. Guo, S. Shrivastava, M. Hassanali, P. Stothard, Z. Chang, J. Woolsey, DrugBank: a comprehensive resource for *In silico* drug discovery and exploration, Nucleic Acids Res. 34 (2006) D668–672. 10.1093/nar/gkj067.

[27] B. Ashraf, N. Atiq, K. Khan, A. Wadood, R. Uddin, Subtractive genomics profiling for potential drug targets identification against *Moraxella catarrhalis*, PLoS One. 17 (2022) e0273252. 10.1371/journal.pone.0273252.

[28] E. Gasteiger, A. Gattiker, C. Hoogland, I. Ivanyi, R.D. Appel, A. Bairoch, ExPASy: The proteomics server for in-depth protein knowledge and analysis, Nucleic Acids Res. 31 (2003) 3784–3788. 10.1093/nar/gkg563.

[29] C. Geourjon, G. Deleage, SOPMA: significant improvements in protein secondary structure prediction by consensus prediction from multiple alignments, Comput Appl Biosci. 11 (1995) 681–684. 10.1093/bioinformatics/11.6.681.

[30] T. Schwede, J. Kopp, N. Guex, M.C. Peitsch, SWISS-MODEL: An automated protein homology-modeling server, Nucleic Acids Res. 31 (2003) 3381–3385. 10.1093/nar/gkg520.

[31] R.A. Laskowski, M.W. MacArthur, D.S. Moss, J.M. Thornton, PROCHECK: a program to check the stereochemical quality of protein structures, Journal of Applied Crystallography. 26 (1993) 283–291. doi:10.1107/S0021889892009944.

[32] A. Volkamer, D. Kuhn, F. Rippmann, M. Rarey, DoGSiteScorer: a web server for automatic binding site prediction, analysis and druggability assessment, Bioinformatics. 28 (2012) 2074–2075. 10.1093/bioinformatics/bts310.

[33] E.C. Meng, E.F. Pettersen, G.S. Couch, C.C. Huang, T.E. Ferrin, Tools for integrated sequence-structure analysis with UCSF Chimera, BMC Bioinformatics. 7 (2006) 339. 10.1186/1471-2105-7-339.

[34] S.S. Costa, L.C. Guimaraes, A. Silva, S.C. Soares, R.A. Barauna, First Steps in the Analysis of Prokaryotic Pan-Genomes, Bioinform Biol Insights. 14 (2020) 1177932220938064. 10.1177/1177932220938064.

[35] R.C. Inglin, L. Meile, M.J.A. Stevens, Clustering of Pan- and Core-genome of *Lactobacillus* provides Novel Evolutionary Insights for Differentiation, BMC Genomics. 19 (2018) 284. 10.1186/s12864-018-4601-5.

[36] L.A. Gallagher, S.A. Lee, C. Manoil, Importance of Core Genome Functions for an Extreme Antibiotic Resistance Trait, mBio. 8 (2017). 10.1128/mBio.01655-17.

[37] T. Hossain, M. Kamruzzaman, T.Z. Choudhury, H.N. Mahmood, A. Nabi, M.I. Hosen, Application of the Subtractive Genomics and Molecular Docking Analysis for the Identification of Novel Putative Drug Targets against *Salmonella enterica* subsp. *enterica serovar* Poona, Biomed Res Int. 2017 (2017) 3783714. 10.1155/2017/3783714.

[38] A.M. Butt, I. Nasrullah, S. Tahir, Y. Tong, Comparative genomics analysis of *Mycobacterium ulcerans* for the identification of putative essential genes and therapeutic candidates, PLoS One. 7 (2012) e43080. 10.1371/journal.pone.0043080.

[39] F. Yan, F. Gao, A systematic strategy for the investigation of vaccines and drugs targeting bacteria, Comput Struct Biotechnol J. 18 (2020) 1525–1538. 10.1016/j.csbj.2020.06.008.

[40] A.R. Oany, M. Mia, T. Pervin, M.N. Hasan, A. Hirashima, Identification of potential drug targets and inhibitor of the pathogenic bacteria *Shigella flexneri* 2a through the subtractive genomic approach, In silico Pharmacol. 6 (2018) 11. 10.1007/s40203-018-0048-2.

[41] A.J. Fatoba, M. Okpeku, M.A. Adeleke, Subtractive Genomics Approach for Identification of Novel Therapeutic Drug Targets in *Mycoplasma genitalium*, Pathogens. 10 (2021). 10.3390/pathogens10080921.

[42] M. Duffield, I. Cooper, E. McAlister, M. Bayliss, D. Ford, P. Oyston, Predicting conserved essential genes in bacteria: *In silico* identification of putative drug targets, Mol Biosyst. 6 (2010) 2482–2489. 10.1039/c0mb00001a.

[43] K. Khan, K. Jalal, A. Khan, A. Al-Harrasi, R. Uddin, Comparative Metabolic Pathways Analysis and Subtractive Genomics Profiling to Prioritize Potential Drug Targets Against *Streptococcus pneumoniae*, Front Microbiol. 12 (2021) 796363. 10.3389/fmicb.2021.796363.

[44] R. Gupta, R. Verma, D. Pradhan, A.K. Jain, A. Umamaheswari, C.S. Rai, An *In silico* approach towards identification of novel drug targets in pathogenic species of *Leptospira*, PLoS One. 14 (2019) e0221446. 10.1371/journal.pone.0221446.

[45] D.H. Kim, Q.T. Nguyen, G.S. Ko, J.K. Yang, Molecular and Enzymatic Features of Homoserine Dehydrogenase from *Bacillus subtilis*, J Microbiol Biotechnol. 30 (2020) 1905–1911. 10.4014/jmb.2004.04060.

[46] L. Hor, R.C. Dobson, M.T. Downton, J. Wagner, C.A. Hutton, M.A. Perugini, Dimerization of bacterial diaminopimelate epimerase is essential for catalysis, J Biol Chem. 288 (2013) 9238–9248. 10.1074/jbc.M113.450148.

[47] W. Min, H. Li, H. Li, C. Liu, J. Liu, Characterization of Aspartate Kinase from *Corynebacterium pekinense* and the Critical Site of Arg169, Int J Mol Sci. 16 (2015) 28270–28284. 10.3390/ijms161226098.

[48] T.C. Ullrich, M. Blaesse, R. Huber, Crystal structure of ATP sulfurylase from *Saccharomyces cerevisiae*, a key enzyme in sulfate activation, EMBO J. 20 (2001) 316–329. 10.1093/emboj/20.3.316.

[49] A. Bairoch, The ENZYME database in 2000, Nucleic Acids Res. 28 (2000) 304–305. 10.1093/nar/28.1.304.

[50] A. Stenlund, Initiation of DNA replication: lessons from viral initiator proteins, Nat Rev Mol Cell Biol. 4 (2003) 777–785. 10.1038/nrm1226.

[51] C. Yu, F. Yang, D. Xue, X. Wang, H. Chen, The Regulatory Functions of sigma(54) Factor in Phytopathogenic Bacteria, Int J Mol Sci. 22 (2021). 10.3390/ijms222312692.

[52] C. Ingram-Smith, J. Wharton, C. Reinholz, T. Doucet, R. Hesler, K. Smith, The Role of Active Site Residues in ATP Binding and Catalysis in the *Methanosarcina thermophila* Acetate Kinase, Life (Basel). 5 (2015) 861–871. 10.3390/life5010861.

[53] H. Ritz, N. Schramek, A. Bracher, S. Herz, W. Eisenreich, G. Richter, A. Bacher, Biosynthesis of riboflavin: studies on the mechanism of GTP cyclohydrolase II, J Biol Chem. 276 (2001) 22273–22277. 10.1074/jbc.M100752200.

[54] J.K. Smith, J.V. Schloss, B.J. Mazur, Functional expression of plant acetolactate synthase genes in *Escherichia coli*, Proc Natl Acad Sci U S A. 86 (1989) 4179–4183. 10.1073/pnas.86.11.4179.

[55] A. Economou, Bacterial preprotein translocase: mechanism and conformational dynamics of a processive enzyme, Mol Microbiol. 27 (1998) 511–518. 10.1046/j.1365-2958.1998.00713.x.

[56] U. Hafsa, G.S. Chuwdhury, M.K. Hasan, T. Ahsan, M.A. Moni, An *In silico* approach towards identification of novel drug targets in *Klebsiella oxytoca*, Informatics in Medicine Unlocked. 31 (2022) 100998. 10.1016/j.imu.2022.100998.

[57] S.S. Hassan, R. Shams, I. Camps, Z. Basharat, S. Sohail, Y. Khan, A. Ullah, M. Irfan, J. Ali, M. Bilal, C.M. Morel, Subtractive sequence analysis aided druggable targets mining in *Burkholderia cepacia* complex and finding inhibitors through bioinformatics approach, Mol Divers. 27 (2023) 2823–2847. 10.1007/s11030-022-10584-5.

[58] K. Ogata, Y. Yajima, S. Nakamura, R. Kaneko, M. Goto, T. Ohshima, K. Yoshimune, Inhibition of homoserine dehydrogenase by formation of a cysteine-NAD covalent complex, Sci Rep. 8 (2018) 5749. 10.1038/s41598-018-24063-1.

[59] F. Schobel, I.D. Jacobsen, M. Brock, Evaluation of lysine biosynthesis as an antifungal drug target: biochemical characterization of *Aspergillus fumigatus* homocitrate synthase and virulence studies, Eukaryot Cell. 9 (2010) 878–893. 10.1128/EC.00020-10.

[60] A. Garg, R. Tewari, G.P. Raghava, Virtual Screening of potential drug-like inhibitors against Lysine/DAP pathway of *Mycobacterium tuberculosis*, BMC Bioinformatics. 11 Suppl 1 (2010) S53. 10.1186/1471-2105-11-S1-S53.

[61] S. Muduli, S. Karmakar, S. Mishra, The coordinated action of the enzymes in the L-lysine biosynthetic pathway and how to inhibit it for antibiotic targets, Biochim Biophys Acta Gen Subj. 1867 (2023) 130320. 10.1016/j.bbagen.2023.130320.

